# Potential Achilles heels of SARS-CoV-2 are best displayed by the base order-dependent component of RNA folding energy

**DOI:** 10.1101/2020.10.22.343673

**Authors:** Chiyu Zhang, Donald R. Forsdyke

## Abstract

The base order-dependent component of folding energy has revealed a highly conserved region in HIV-1 genomes that associates with RNA structure. This corresponds to a packaging signal that is recognized by the nucleocapsid domain of the Gag polyprotein. Long viewed as a potential HIV-1 “Achilles heel,” the signal can be targeted by a new antiviral compound. Although SARS-CoV-2 differs in many respects from HIV-1, the same technology displays regions with a high base order-dependent folding energy component, which are also highly conserved. This indicates structural invariance (SI) sustained by natural selection. While the regions are often also protein-encoding (e.g. NSP3, ORF3a), we suggest that their nucleic acid level functions can be considered potential “Achilles heels” for SARS-CoV-2, perhaps susceptible to therapies like those envisaged for AIDS. The ribosomal frameshifting element scored well, but higher SI scores were obtained in other regions, including those encoding NSP13 and the nucleocapsid (N) protein.

## 1. Introduction

After four decades of research, infection with HIV-1 can routinely be controlled but, because the virus adopts DNA-form latency, not cured. Since the newly emergent SARS-CoV-2 displays neither HIV-1-like latency nor its extreme variability and chronicity, curative treatments would seem more feasible (Rausch et al., 2020). Although both are RNA viruses, their many differences indicate that different therapeutic strategies will be required. Yet, as with HIV-1, strategies for SARS-CoV-2 have focused, with limited success, on the functions of encoded proteins rather than those of its RNA genome or of transcripts from that genome. Intriguingly, a nucleic acid level bioinformatics technology that long ago suggested a HIV-1 vulnerability related to genome function (Forsdyke 1995a, 2014), has recently gained support (Ingemarsdotter et al., 2018; Rein 2020; Ding et al., 2020), and the term “Achilles heel” is increasingly employed (Ding et al., 2020; Forsdyke 2016). The same technology now suggests similar vulnerabilities for SARS-CoV-2 (Rausch et al., 2020).

Our analytical approach requires elimination of the contribution of base composition to the energetics of folding into a higher order (stem-loop) structure of a single-stranded nucleic acid. Just as accent or dialect affects a spoken text in its entirety, so base composition tends to reflect *genome-wide* evolutionary pressures. Just as a local arrangement of words best conveys specific meaning to a text, so base order best reflects *local* evolutionary pressures. Base order is most likely to be conserved when encoding a function critical for survival. Because they do not make this fundamental distinction between base composition and order, more sophisticated methods of SARS-CoV-2 structure determination that incorporate all sequence information (Huston et al., 2020; Tavares et al., 2020; Manfredonia et al., 2020), have tended to confuse rather than clarify. Our relatively simple assays of the base order-dependent component of the folding energy, which are devoid of redundant base compositional information (see Materials and Methods), have shown that a highly conserved region, in otherwise rapidly mutating HIV-1 genomes, associates with an RNA structure corresponding, not to a protein-encoding function, but to an RNA packaging signal. The latter is specifically recognized by the nucleocapsid domain of the Gag polyprotein (Sarni et al., 2020) and is now seen as a potential “Achilles heel” of HIV-1 that can be targeted by a recently described antiviral compound (Ingemarsdotter et al., 2018).

We here report similar highly conserved structural regions of the SARS-CoV-2 genome, one or more of which should be susceptible to targeting (Medeiros et al., 2020; Haniff et al., 2020). We identify certain open reading frames (ORFs) that, because of their conservation, have so far attracted therapeutic interest mainly related to their functions at the protein level (Robson 2020; Issa et al., 2020), rather than at the level of the corresponding, yet highly structured, regions of the genome. The ribosomal frameshift element (FSE) that is among our results, is attracting attention (Haniff et al., 2020; Kelly et al., 2020; Ziv et al., 2020). Yet our analytical approach, which is employed by others (Andrews et al. 2018, 2020; Simmonds 2020a, b), suggests there may be more suitable targets in other regions.

## 2. Materials and methods

### 2.1. Sequences

These were obtained from the NCBI (Bethesda) and GISAID EpiCoV (Munich) databases. The Wuhan-Hu1 sequence (GenBank NC_045512.2), deemed taxonomically prototypic (Kumar et al., 2020; Pipes et al., 2020), was compared (regarding base substitutions and folding potential) with 381 Chinese isolates, 430 Italian isolates, and 932 isolates from New York, USA. Our “window” starting point was base 1 of the 29,903 base prototype sequence. We refer to windows by their centers. The center of the first 200 base window would be 100.

### 2.2. Substitution frequencies

In previous HIV-1 studies the base differences between just two individual sequences sufficed for the tabulation of a statistically significant set of base substitution frequencies (Forsdyke 1995a). The lower mutation rates of SARS-CoV-2 strains (Robson 2020) required the tabulation of changeable positions relative to the prototype (i.e. polymorphisms) among a set of individual sequences in a geographical region. In early September (2020), 709 Chinese sequences were filtered to remove duplicates and those with ambiguities, so yielding 381 unique sequences. These were scored for substitutable positions (whether one base could be exchanged by one of the three alternative bases) by aligning against the Wuhan prototype using Muscle software implemented in MEGA 7.0 (Edgar 2004), with manual adjustments. This yielded a substitution value for the 200 bases in each sequence window (i.e. values ranging from zero to 200). The values were in the low range indicating less likelihood of back mutations (site saturation) in the period since divergence from the presumed prototype. Indeed, in a study of several thousand genomes 73% of sites were conserved (i.e. only 27% were polymorphic) (Simmonds 2020b). To gain support for our results, in late September our study was extended with sets from Italy (430 from 642 downloaded) and New York, USA (932 out of 1483 downloaded). As expected, the FORS-D profiles of two early-arising members of these sets, that differed from the prototype in two (Italy) and nine (USA) bases, scarcely varied from the profile of the Chinese prototype shown here (i.e. overlapping profiles). In contrast, profiles differ greatly when different coronavirus species are compared (Simmonds 2020b).

### 2.3. Base order-dependent component of folding energy

The energetics of the folding of a single-stranded nucleic acid into a stem-loop structure depend on both the composition and order of its bases. Base composition is a distinctive characteristic of a genome or large genome sector. A localized sequence (e.g. a 200 base window), which is rich in the strongly-pairing bases G and C, will tend to have a stable structure simply by virtue of its base composition, rather than of its unique base order. This high GC% value can obscure the contribution of the base order-dependent component of the folding energy, which provides a sensitive indicator of *local intraspecies* pressures for the conservation of function within a population (i. e. a mutated organism is eliminated by natural selection so no longer can be assayed for function in the population). In contrast, *interspecies* mutations tend to influence *genome-wide* oligonucleotide (k-mer) pressure, of which base composition (GC%) is an indicator (Aggarwala and Voight 2016; Morozov 2017). Assays of the pressure can assist the initial stages of a formal sequence alignment, but this must then be finalized by attending to local details (Edgar 2004). Oligonucleotide (k-mer) pressure works to generate and/or sustain members of emerging species by preventing recombination with parental forms (Forsdyke 1996, 2014, 2016, 2019a, b). Its elimination facilitates focus on local folding.

Early studies of RNA virus structure by Le and Maizel (1989) were primarily concerned with the statistical significance of RNA folding, rather than with distinguishing the relative contributions of base composition and base order. However, with a pipeline between the various programs that were offered by the Wisconsin Genetics Computer Group, the base composition and base order-dependent components were separated and individually assessed (“folding of randomized sequence difference” analysis; FORS-D analysis). Departing from Le and Maizel, FORS-D values (see below) were not divided to yield Z-scores, but were simply plotted with statistical error bars (Forsdyke 1995a,b,c, 2013, 2014, 2016). The limits of the latter were generally close to the corresponding FORS-D values and, for clarity, are omitted here.

A window of 200 bases is moved along a natural sequence in 20 or 50 base steps. A folding program (Zuker 1989) is applied to the sequence in each window to obtain “folding of natural sequence” (FONS) values for each window, to which both base composition and base order will have contributed. The four bases in each sequence window are then shuffled to destroy their order while retaining base composition, and folding energy is again determined. This shuffle-and-fold “Monte Carlo” procedure is repeated ten times and the average (mean) folding value is taken as the “folding of randomized sequence mean” (FORS-M) value for that window. This reflects the contribution of base composition alone. The base order-dependent component is then derived by subtraction from the FONS value. This is the “folding of randomized sequence difference” (FORS-D) value. The sign of the difference value depends on the direction of the subtraction. (FORS-D was given positive values in the early 1990s, but the direction of subtraction was changed in subsequent work.) Fluctuations seen in FONS profiles of genomes are mostly due to changes in the FORS-D component, whereas the FORS-M component, while making a major contribution to folding energetics, is relatively constant.

### 2.4. Validation of four base shuffling

The approach was modified by others who, rather than shuffling the *four* bases, favored retaining some base order information. Accordingly, they shuffled groups of bases (e.g. the *sixteen* dinucleotides). Following disparagement of the conceptual basis of four base shuffling, which was duly clarified (Forsdyke 2007a), the validity of single base level shuffling is now generally accepted and is being applied routinely to viral genomes (Andrews et al. 2018, 2020; Simmonds 2020a, b). The Monte Carlo procedure can also be simplified to decrease FORS-M computational time (Chen et al., 1990) using support vector machine-based technology (Washietl et al. 2005). A later software modification (“Bodslp”), written by Professor Jian-sheng Wu (Zhang et al., 2008), retains our Monte Carlo approach and was further developed by Professor Shungao Xu as “Random Fold-Scan” for Windows-based systems (Xu et al., 2007).

In addition to assisting the study of infectious viruses and protozoa (Xue and Forsdyke 2003), FORS-D analysis proved fruitful when applied to topics such as speciation (Forsdyke 1996, 2014; Zhang et al., 2008), the origin of introns (Forsdyke 1995b, 1995c, 2013), relating structure to recombination breakpoints and deletions (Zhang et al., 2005a, b), and relying on a single sequence (rather than alignments) for the determination of positive Darwinian selection (Forsdyke 2007b).

However, for a given sequence window, output can follow only from the base order *in that window*. Lost are higher order structures that might occur naturally through long-range interactions (Ziv et al., 2020). Furthermore, if the artificial demarcation of a window happens to cut between the two limbs of a natural stem-loop structure, then lost are what might have been contributed to the folding energetics had a larger window, or a different section point, been chosen. Variations in step size will generate differences in windows and hence variations in results. Such variations might be less when window margins correspond to natural section points, such as those demarcating an RNA transcript. There is also a kinetic aspect, particularly apparent with transcripts, due to the probability that the natural pattern of early 5′ folding will influence later folding.

### 2.5. Validation of structural invariance index

Plots of structural invariance (SI) facilitate the identification of conserved RNA structures. Structure and invariance can be united numerically by addition (SI = FORS-D + Substitutions). Structure is reflected in the degree of *negativity* of FORS-D values, and invariance is reflected in limitation of recorded *positive* substitutions at individual base sites (polymorphisms). Thus, the SI would be −20 units in the case of a FORS-D value of −20 kcal/mol in a window where there had been zero recorded substitutions (i.e. mutant forms had been eliminated by natural selection or genetic drift). If there had been 10 recorded substitutions the SI would be −10 units.

High negative SI values indicate structural conservation among a set of genomes. The validity of this approach is supported by prior work with HIV-1. Fig. 1 shows application of the index to previously reported data on HIV-1 (Forsdyke 1995a, 2014). Here a high negative SI index value corresponds to regions recognized as likely to offer Achilles heel-like vulnerability (Forsdyke 2016). The region around the window centered on 500 nucleotide bases is the focus of recent work (Ingemarsdotter et al., 2018; Rein 2020; Ding et al., 2020).

**Fig. 1.**
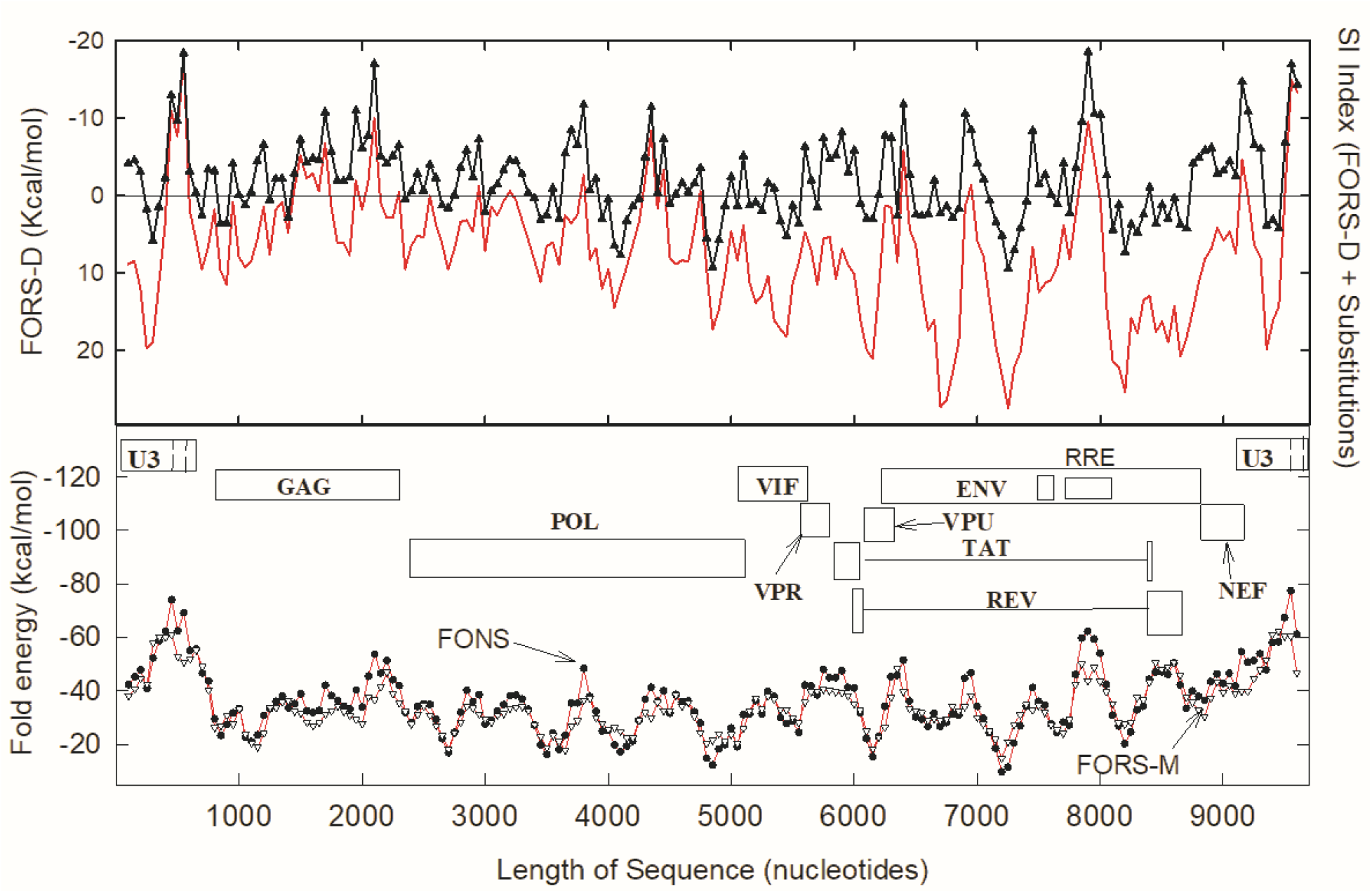
Structural invariance (SI) of HIV-1 (subtype SF2) as compared with reference subtype (HXB2). The region recently recognized as a potential Achilles heel (bases 380-640), has a high negative FORS-D value (black triangles) similar to those of the RRE (Rev response element) and the 3′ untranslated region (UTR). The SI index (continuous red line) indicates highest sequence conservation in the regions of the RNA packaging signal and the 3′UTR. For details see Forsdyke (1995a).

## 3. Results

### 3.1. Variation due to base order component

A nucleic acid folding program was applied to 200 nucleotide base “windows” that were moved along a SARS-CoV-2 sequence in 20 base steps. This provided a set of folding energy values fluctuating from −26 kCal/mol to −78 kCal/mol, with the latter negative value indicating *stronger* folding (Fig. 2; blue top line). These “folding of natural sequence” (FONS) values were further dissected into their two fundamental components, that due to base *composition* (FORS-M) and that due to base *order* (FORS-D).

**Fig. 2.**
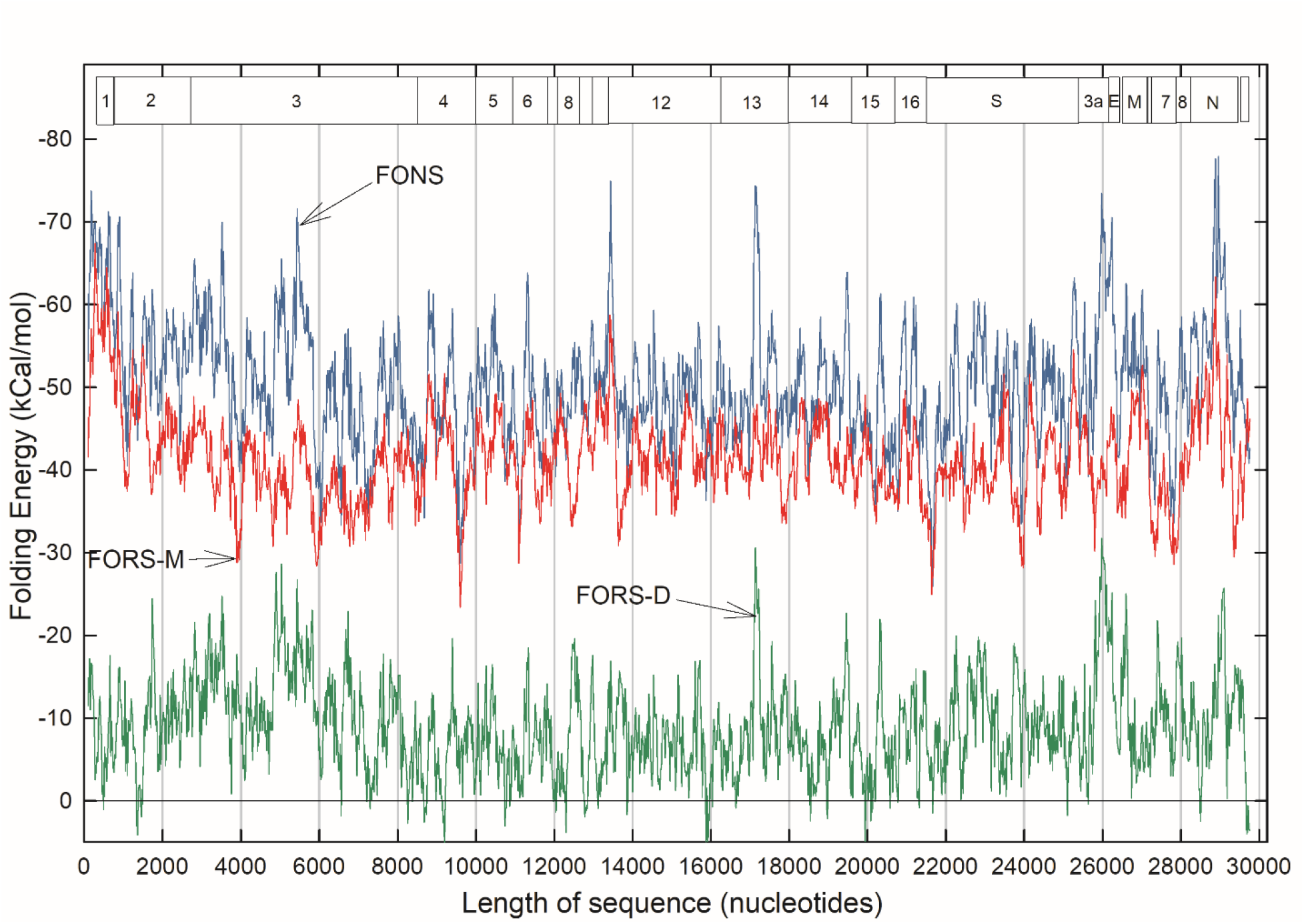
Variations in folding energetics throughout the single-stranded RNA genome of a clade B SARS-CoV-2 sequence, deemed “prototypic” since it appeared in China early in the COVID-19 epidemic (Wuhan-Hu1; NC_045512.2). Data points for 1484 200 base windows (beginning at base 1 and moved in *20 base steps*) were connected rectilinearly. FONS values (top, blue); FORS-M values (middle, red); FORS-D values (bottom, green). Boxes indicate the locations of the nonstructural proteins (NSP 1-16) and structural proteins (S, etc.).

The composition component (Fig. 2; red middle line) is generally less negative than the FONS value (blue line), the difference being due to the contribution made by the base order-dependent component (green bottom line). Composition (FORS-M) usually makes a larger, but *less variable* contribution, whereas the order component (FORS-D) usually makes a smaller, but *highly variable* contribution. Thus, much of the fluctuation that distinguishes different parts of a genome in the FONS profile is due to the base order component. As with our previous studies of a wide variety of genomes, FORS-D values provide a sensitive indicator of regional differences in the strength of folding. A notable exception is a peak (−75 kCal/mol) located in a window centered on base 13440 at the boundary between ORFs NSP10 and NSP12 in the region of the ribosomal FSE. Here, the base composition component makes a major contribution.

In a few regions FORS-M is more negative than FONS and here FORS-D values are positive. Thus, here “nature” appears to have arranged the order of the nucleotide bases to *restrain* the impact of base composition on the strength of folding (i.e. encourage formation of loops that might be targeted by antisense RNAs; Forsdyke 2007a). A need for this appears particularly evident in windows centered on the following bases: 1360 (NSP2), 9200 (NSP4), 12300 (NSP8), 15880 and 15920 (NSP12), 19940 (NSP15), 29680 and 29700 (3′ untranslated region).

### 3.2. Inverse relationship between folding and substitutions

The degree of conservation in sequential windows of members of a set of sequences from China was evaluated as the number of substitutable base positions (polymorphism), relative to the prototype sequence (Fig. 3). This was compared with corresponding FORS-D values, the profile of which differed a little from that of Fig. 2 due to different step values (see Materials and Methods).

**Fig. 3.**
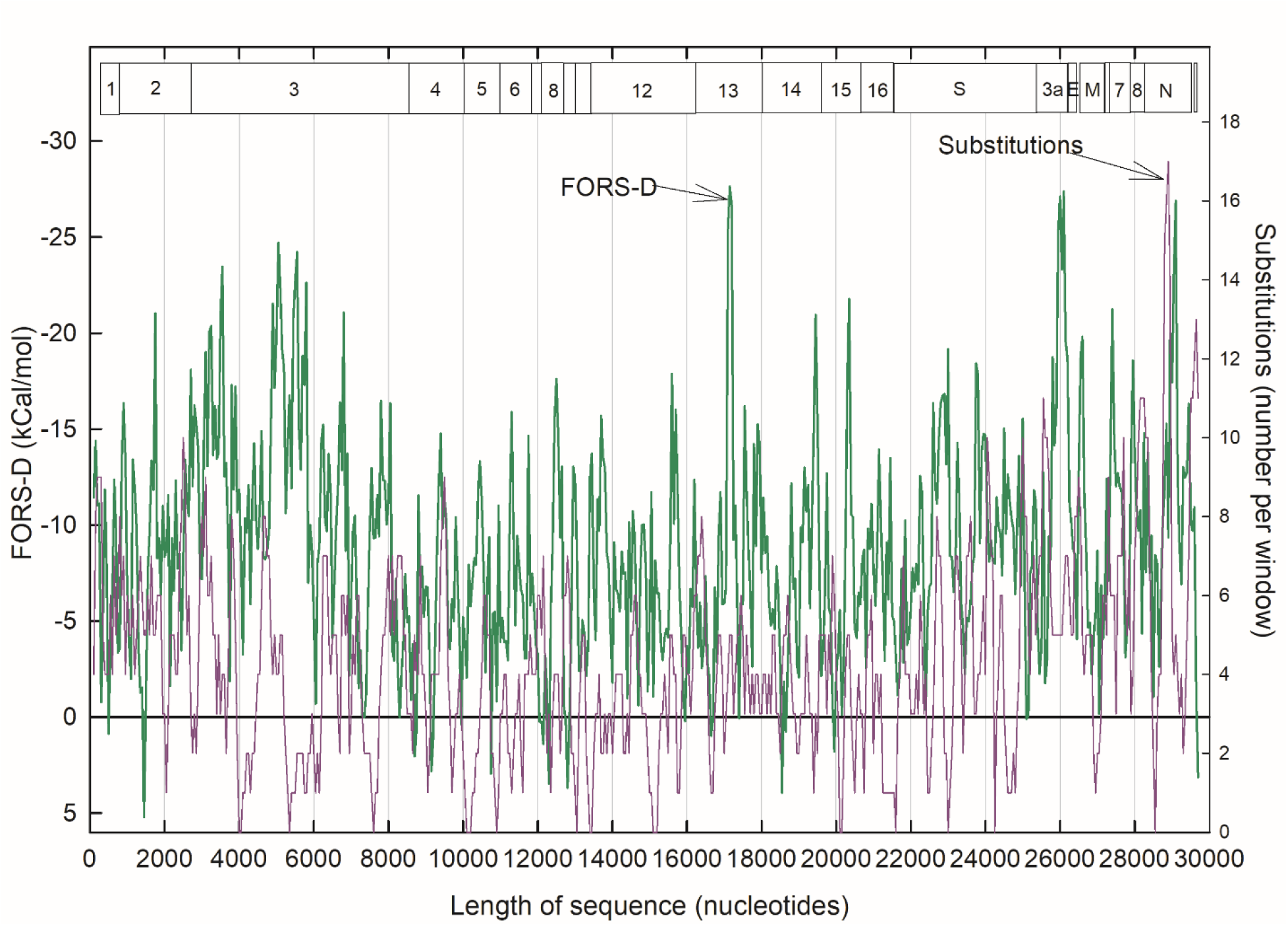
Comparison of base order-dependent stem-loop potential values (FORS-D; green) for the Wuhan-Hu1 SARS-CoV-2 prototype, and base substitutions values (purple). The latter were found in the same sequence windows among a set of Chinese SARS-CoV-2 isolates. Data points for 594 200 base sequence windows (beginning at base 1 and moved in *50 base steps*) were connected rectilinearly. (There were some ambiguities in substitutions in the sequences included in the first two windows, making the first two data points less reliable).

As with previous studies (Forsdyke 1995a, b, c), there tended to be a reciprocal relationship between the base order contribution to folding energy (FORS-D) and substitutions. When one was high the other was low, and vice-versa. Negative FORS-D values were generally greatest in the region of ORF NSP3, and in narrow regions corresponding to the NSP13, ORF3a and the nucleocapsid (N) proteins. Zero substitutions were observed for windows centered on the following base nucleotide values, some isolated and some grouped (in brackets): [4000, 4050], 5350, 7600, [10100, 10150, 10200], 10900, 13050, [13400, 13450; close to ribosomal FSE], [15100, 15150, 15200], [20100, 20150], 21600, 23000, 24250, 28550. Substitutions were higher in the last part of the sequence, being particularly evident in the nucleocapsid (N) protein ORF. While the zero-substitution windows centered on bases 5350, 7600 and 21600 appeared isolated, each had many neighboring windows with few substitutions. At some positions, *low* substitution values were accompanied by *high* negative RNA folding energy values (i.e. reciprocal relationships), indicating conservation of structural function that could be at the RNA genome level or in RNA transcripts.

### 3.3. Structural invariance index identifies potential targets

For a more focused view of the reciprocal separation of high *negative* folding values and corresponding numbers of substitutions (ranging from zero to high *positive* values) the two were added to provide the “structural invariance” (SI) index (Fig. 4).

**Fig. 4.**
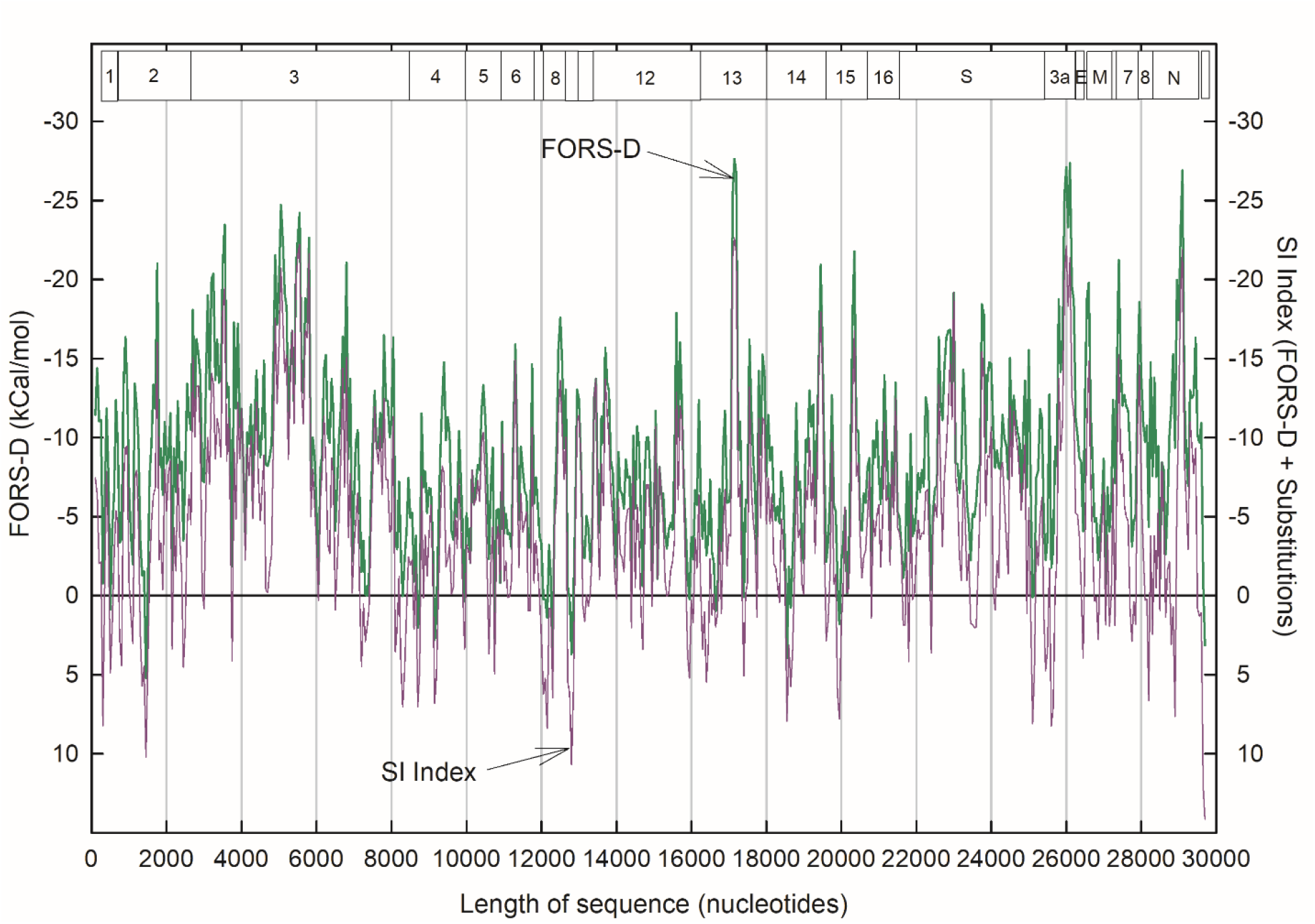
Data of figure 3 plotted with FORS-D values (green) accompanied by the structural invariance (SI) index (purple). (For plots of analogous HIV-1 data see Fig. 1).

Here, despite having some substitutions, ORFs NSP13 and NSP3 were preeminent (scoring – 22.6 and −22.2 SI units, respectively). The next two best scores were within ORF3a (scoring – 22.1 SI units) and within the N protein ORF (scoring −21.9 units). Whereas the scores for the NSP13, ORF3a and N locations were narrowly based, the entire NSP3 location tended towards high scores. Close downstream to the high-scoring N region (window centered on base 29100) is a region with the highest number of substitutions (see Fig. 3) centered on base 28900 (scoring +7.6 SI units).

Windows 13400 and 13450 in the FSE region scored −12.5 and −13.7 units, respectively. However, there were many more windows with higher negative scores. Indeed, SI values more negative than −15.0 units were observed for windows centered on the following base nucleotide values, some isolated and some grouped (in brackets): 1750, 2700, [3500, 3550], 4900, [5000, 5050, 5100], 5200, [5450, 5500, 5550], 5800, 6800, [17100, 17150, 17200], 19450, 20350, 23000, 23750, [25950, 26000, 26050, 26100], 27400, [29050, 29100].

### 3.4. SI indices for Italy and New York (USA)

The SI profile for China (Fig. 4) was, in broad outline, confirmed with corresponding data later downloaded from Italy and New York, USA (Fig. 5). The high *negative* SI indices found in the NSP3, NSP13, ORF3a and N regions, were evident with sequences for all three locations. Other regions, notably the S region, were also corroborated. The high *positive* SI values, indicating regions likely to have poorly conserved structures, are also corroborated at some locations. A high *intraspecies* mutation rate for the N protein ORF (Fig. 3) is also seen when *interspecies* comparisons are made with other coronavirus species, with implications for early speciation mechanisms (Dilucca et al., 2020).

**Fig. 5.**
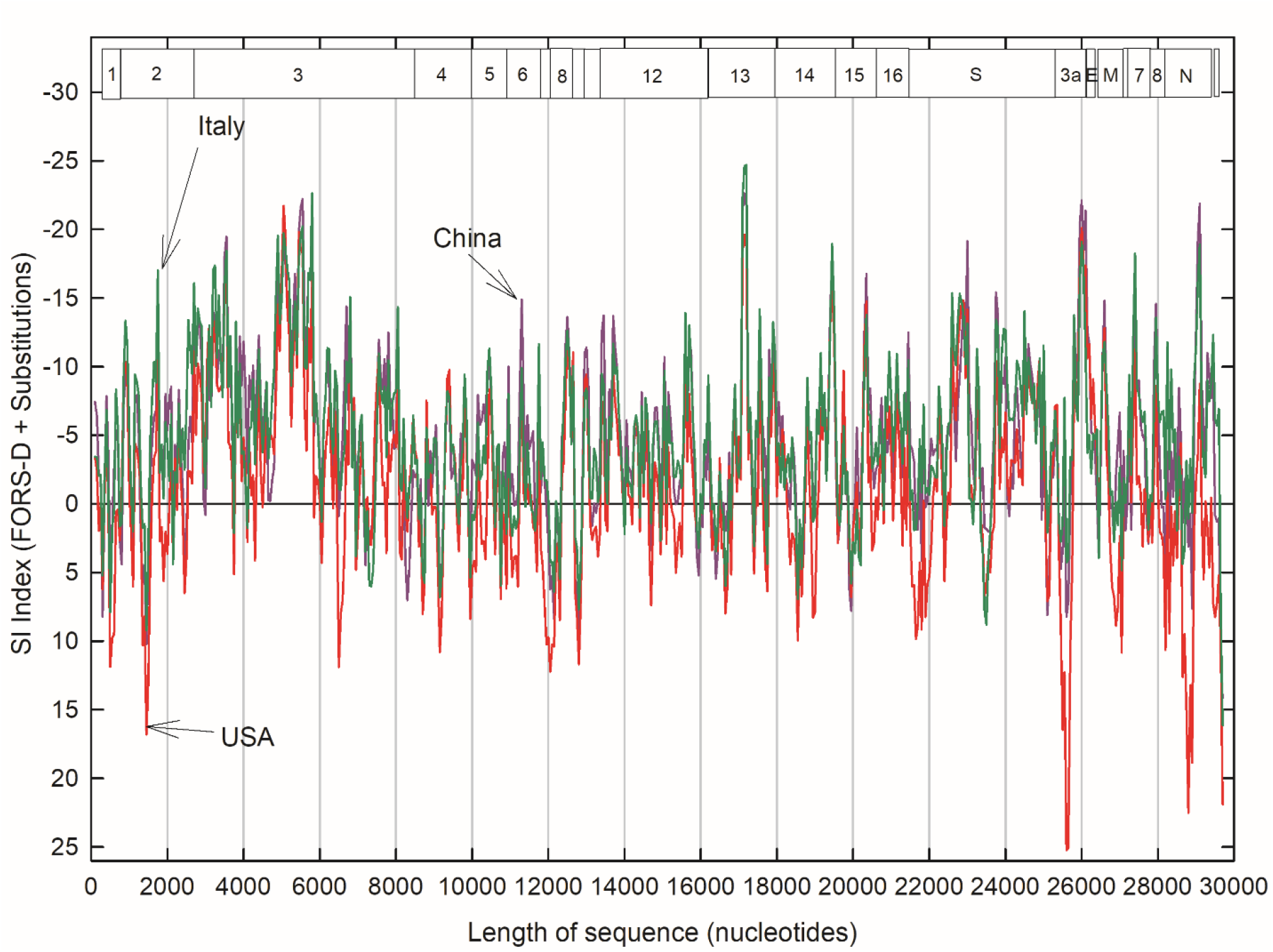
Comparison of SI indices for China (381 isolates) as in Fig. 4 (purple), with SI indices for Italy (430 isolates; green) and New York, USA (932 isolates; red).

## 4. Discussion

### 4.1. Three therapeutic challenges

Mutation rates of microbial pathogens are generally higher than those of their hosts. While a microbe spreading from host-to-host can “anticipate” that it will face a succession of broadly similar challenges, in the short-term those hosts cannot likewise “anticipate” that new microbial invaders will remain as they were in previous hosts. Thus, host immune defenses may be overwhelmed. (In the long-term there is a different scenario related to innate immunity; see below). Therapeutic challenges are, first, to locate a conserved, less-variable, part of a pathogen’s genome that it will have inherited sequentially from a multiplicity of past generations and so is likely to carry through to a multiplicity of future generations. Second, is to identify the corresponding primary function, be it at the genome, RNA transcript, or protein level. Third, from this knowledge (that may be incomplete; i.e. function not fully clarified), devise effective pathogen inhibition without imposing deleterious side-effects on the host.

### 4.2. Identification of potential RNA-level Achilles heels in HIV-1 and SARS-CoV-2

Viral vulnerability is often assumed to associate with protein-level functions (Haniff et al., 2020; Robson 2020). However, studies of the AIDS virus have identified genome structure itself as both functional and conserved, so signifying vulnerability (Forsdyke 1995a, 2014; see Fig. 1). A genomic packaging signal for HIV-1, which is specifically recognized by the nucleocapsid domain of its Gag polyprotein, has long been recognized as a potential “Achilles heel” (Forsdyke, 1995a, 2014, 2016), so inviting therapeutic exploration (Ingemarsdotter et al., 2018; Rein 2020; Ding et al., 2020). Through targeting of specific RNA conformations, Gag not only influences the assembly of HIV-1 genomic RNA into virus particles, but also regulates HIV-1 mRNA translation (Anderson and Lever 2006). RNA conformational flexibility, facilitated by fine-tuned intermediate-strength binding, should permit easier switching between regulatory options. Indeed, small changes in target RNA structure can impede this (Ding et al., 2020). Thus, unlike most other regions of the HIV-1 genome, mutations here would likely lead to negative Darwinian selection at an early stage – hence the high conservation. Application of the same bioinformatic technology to the SARS-CoV-2 virus genome has now revealed similar potential “Achilles heels.”

Lacking the chronicity of HIV-1 infection, the genome of SARS-CoV-2 should have been shaped less by adaptations to counter long-term host immune defenses. It cannot hide within its host genome in latent DNA form. Yet, the larger SARS-CoV-2 genome contains many more genes than HIV, which require differential expression according to the stage of infection. Even more complex regulatory controls can be envisaged, likely requiring conserved genome conformations at appropriate locations. Be they synonymous or non-synonymous, mutations in these structured regions could result in negative selection of the viruses in which they occurred – hence high conservation. The ribosome FSE located close to base 13468 (Figs. 2) would seem to exemplify this (Kelly et al., 2020), and a potential targeting agent is now available (Haniff et al., 2020). However, we have here identified other structurally important regions with more base order-dependence and higher degrees of conservation (Figs 3–5), that might, either singly or collectively, be better candidates for targeting. These broad regions, demarcated by window boundaries, should serve to focus attention on the local structural details required, not only for therapeutic purposes, but also to guide the choice of primers in diagnostic PCR assays.

### 4.3. Base composition relates to species evolution

When determining folding energy, our approach depends on eliminating contributions of base composition which, as noted, plays an unusual role in the case of the FSE. More usually, base composition is a distinctive characteristic of *entire* genomes or large genome sectors, which reflects their underlying oligomer (“k-mer”) content (Aggarwala and Voight 2016; Morozov 2017). The slow genome-wide accumulation of mutations in oligomer composition, easiest documented as changes in 1-mer frequency (base composition; GC%), can serve to initiate divergence into new species. By preventing that accumulation, potentially diverging organisms can stay within the confines of their species (Forsdyke 1996, 2019a, 2019b). The presence or absence of synonymous mutations (Simmonds 2020a, b; Wang et al., 2021), which affect structure rather than amino acid composition, can have an important role in this process. The primary role of *constancy* in the base composition-related character is to prevent recombination with allied species (*interspecies* recombination) while facilitating the *intraspecies* recombination that can correct mutations, so retaining species individuality (Forsdyke 2014, 2016). Such recombination is initiated by “kissing” interactions between complementary sets of unpaired bases (k-mers) at the tips of stem-loop structures (Forsdyke 1996).

Thus, we are here concerned with *localized* intraspecies mutations that affect fitness, so making members of a species carrying those mutations liable to natural selection. The mutations facilitate *within-species* evolution rather than divergence into new species. And when that evolution has run its course, some of the polymorphic bases will have become less mutable, so will be deemed “conserved.” Indeed, mutations of ORF NSP3 are high when the sequences of *different* coronavirus species are compared (Claverie 2020), yet when, from *intraspecies* comparisons, mutations (in the form of base substitutions) are scored, they are very low (Figs. 3, 4). Our technology (see Materials and Methods) removes the base composition-dependent component of mutational changes (that relates more to *interspecies* evolution) and focuses on the base order-dependent component (that relates more to *intraspecies* evolution). It best reflects localized functions, be they encoding protein or determining the potentiality for folding into higher order structure, of linear, single-stranded, nucleic acid sequences.

### 4.4. Conservation as a reliable indicator

We sought regions that were both high in stem-loop potential and bereft of mutations, following the premise that conserved functions would be best targeted therapeutically, assuming the availability of pathogen-specific therapeutic agents that would not cross-react with hosts. Interference with structural nucleic acid level functions might be less likely to produce unforeseen host side-effects than with protein-level functions. But is conservation necessarily a good indicator of likely therapeutic success? Indeed, a conserved function in a pathogen could *owe* that conservation to the pathogen strategy of, whenever possible, mutating to resemble its host. As reviewed elsewhere (Forsdyke 2019c), this would make it less vulnerable to host innate and acquired immune defenses. To prevent autoimmunity, the generation of immune cell repertoires involves the negative selection of self-reacting cells so creating “holes” in the repertoire that pathogens can exploit by progressive mutation towards host-self, testing mutational effectiveness a step at a time. This would make it advantageous for the host, during the process of repertoire generation, not only to *negatively* select immune cells of specificity towards “self,” but also to *positively* select immune cells of specificity towards “near-self.” A high level of anti-near-self immune cell clones would constitute a barrier limiting the extent of pathogen mutation towards self. The existence of such positive selection is now generally accepted, with the implication that some pathogen functions might have approached so close to host-self that targeting them therapeutically would result in cross-reactivity.

This caveat aside, we deem conservation a reliable indicator that a certain pathogen function is likely to be a suitable target for therapy. Attacking a short segment of a pathogen nucleic acid sequence is unlikely to ensnare a similar segment of its host’s nucleic acid. In any case, to militate against this, the pathogen specificity of a potential therapeutic agent can be screened against the prototypic human genomic sequence (assuming it is unlikely that patient genomes will significantly depart from this).

### 4.5. Concluding remarks

While prospects for the development of prophylactic vaccines against infection with SARS-CoV-2 are promising, methods to boost post-infection host immune defenses and to directly target SARS-CoV-2 are urgently needed. These require a better understanding both of viral interactions with host innate and acquired immune systems (Forsdyke 2019c), and of viral vulnerabilities. The latter enquiry proceeds in three steps: Find specific “Achilles heels.” Design therapies to exploit them. Prove their clinical effectiveness. We have here been concerned with the first step – the identification of high SI scoring regions that may then serve to focus attention on local structural details (Wacker et al., 2020; Huston et al., 2020). Although the bioinformatic technology related to this has long been available (Forsdyke 1995a), our claim to reveal viral “Achilles heels,” as promulgated in successive textbook editions (Forsdyke 2016), has only recently gained support (Ingemarsdotter et al., 2018; Rein 2020; Ding et al., 2020). This may be because the importance of removing redundant information and analyzing solely the contribution of base order to folding energy, was not fully appreciated. Even those who have employed the same technology have expressed puzzlement at the “biological purpose” of so much “pervasive RNA secondary structure in the genomes of SARS-CoV-2 and other coronaviruses” (Simmonds 2020a) and regret the “poorly understood RNA structure-mediated effects on innate and adaptive host immune responses” (Simmonds et al., 2020). Thus, we have here repeated and expanded on past clarifications (Forsdyke 2007a; Xu et al., 2007; Zhang et al., 2008) of the conceptual basis of a technology that has contributed to the understanding of a many biological problems other than viral infections (Forsdyke 2016). Meanwhile, it is pleasing to note that, with minimal evidence on targeting, progress is being made with the second step (Medeiros et al., 2020; Haniff et al., 2020; Chen et al. 2020; Suresh et al., 2020). It is hoped that by targeting one or more of the conserved regions in the SARS-CoV-2 genome here identified, rapid cures will be achieved.

## Acknowledgements

We thank Prof. Shungao Xu at Jiangsu University for software, and Ms. Le Cao and Ms. Yingying Ma at Shanghai Public Health Clinical Center, Fudan University, for their technical support. Queen’s University hosts Forsdyke’s webpages. The bioRxiv server hosts a preprint.

